# Acute Thiamethoxam toxicity in honeybees is not enhanced by common fungicide and herbicide and lacks stress-induced changes in mRNA splicing

**DOI:** 10.1101/641407

**Authors:** Pâmela Decio, Pinar Ustaoglu, Thaisa C. Roat, Osmar Malaspina, Jean-Marc Devaud, Reinhard Stöger, Matthias Soller

## Abstract

Securing food supply for a growing population is a major challenge and heavily relies on the use of agrochemicals to maximize crop yield. Neonicotinoids are globally one of the most widely used insecticides. It is increasingly recognized, that some neonicotinoids have a negative impact on non-target organisms, including important pollinators such as the European honeybee *Apis mellifera*. Toxicity of neonicotinoids may be enhanced through simultaneous exposure with additional pesticides, which could help explain, in part, the global decline of honeybee colonies. Here we examined whether exposure effects of the neonicotinoid Thiamethoxam are enhanced by the commonly used fungicide Carbendazim and the herbicide Glyphosate. We also analysed alternative splicing changes upon pesticide exposure in the honeybee. In particular, we examined transcripts of three genes: i) the stress sensor gene *X box binding protein-1* (*Xbp1)*, ii) the *Down Syndrome Cell Adhesion Molecule* (*Dscam)* gene and iii) the *embryonic lethal/abnormal visual system* (*elav)* gene, which are important for neuronal function. Our results showed that acute neonicotinoid exposure is not enhanced by Carbendazim, nor Glyphosate. Toxicity of the compounds did not trigger stress-induced, alternative splicing in the analysed mRNAs, thereby leaving dormant a cellular response pathway to these man-made environmental perturbations.

## INTRODUCTION

Worldwide honeybees and other insects encounter new man-made compounds at potentially harmful concentrations in agricultural landscapes. The combinatorial use of many herbicides, fungicides and pesticides is increasingly recognized for having a negative impact on many pollinators including the honeybee *Apis mellifera* ^1,2^. Forager bees may be exposed to chemicals applied to crops during their foraging for nectar, pollen and water ^3,4^. Through the contaminated food harvested by bees and brought into the hive, the entire colony can be exposed to complex cocktails of xenobiotics ^5^. Such exposure to sub-lethal mixtures of pesticides may cause a reduction in vigour and productivity of the hive ^5,6^. Indeed, honeybee colonies are in decline in many parts of the world and numerous interacting factors are thought to drive the rates of loss, including pathogens, poor nutrition, environmental stress and crop protection chemicals [reviewed in ^7^].

One class of insecticides used globally are the neonicotinoids. These nicotine-like neurotoxic insecticides have been linked to declining bee health ^8^. At high levels, neonicotinoids lead to paralysis and death of target and non-target insects by binding to nicotinic acetylcholine receptors (nAChRs) which are expressed in the insect nervous system ^9,10^. Thiamethoxam is one of the neonicotinoid compounds known to affect honeybees ^11–14^.

Glyphosate is the most widely applied herbicide worldwide ^15^ and often detected in honey, wax, pollen, and nectar ^16–18^. Generally considered harmless to pollinating insects, glyphosate has been reported to affect larval development and feeding behaviour of honeybees ^19,20^. Likewise, the fungicide carbendazim can persist in the environment due to its hydrolytically stable properties ^21,22^ and is a frequent contaminant of bee hives ^23^.

The honeybee genome encodes a comparatively small repertoire of xenobiotic detoxifying enzymes ^24^. Consequently, honeybees have only limited physiological and cellular response options when confronted with different mixtures of agrochemicals. A potential cellular strategy to rapidly respond to such environmental stressors would be differential expression and processing of messenger RNAs (mRNAs) ^25^. Alternative splicing in particular enables cells of an organism to alter and expand availability of different transcripts and their encoding protein-isoforms in response to environmental perturbations ^26,27^. Sub-lethal exposure of xenobiotics can, indeed, alter gene expression ^28^ and induce modulation of splicing reactions ^29,30^. Investigation of potential splicing effects mediated through the action of pesticides may help to clarify how toxic agents interfere with honeybee metabolism.

The *X box binding protein-1* (*Xbp1*) mediates the unfolded protein response (UPR) as a reaction to cellular stress through a splicing mechanism ^31–34, 59^. The *Xbp1* mRNA contains a retained intron that prevents expression of functional Xbp1 protein. This intron is spliced through a mechanism normally operative in tRNA genes leading to expression of the full length Xbp1 transcription factor, which then triggers the UPR.

Alternative mRNA splicing is particularly abundant in the brain and most elaborate in ion channels and cell adhesion molecule genes ^25,35,36^. The most extraordinary example of an alternatively spliced gene is *Down syndrome cell adhesion molecule* (*Dscam*) gene in the fruitfly *Drosophila melanogaster*. *Dscam* can encode 38,016 alternatively spliced mRNAs. *Dscam* plays important roles in neuronal wiring and axon guidance in the nervous system and in phagocytosis of pathogens in the immune system ^37–41^. Moreover, genetic variants of *Dscam* have been linked to insecticide resistance in *Drosophila* ^42^. *Dscam* alternative splicing has been studied in exon-clusters 4, 6 and 9, which harbour an array of mutually exclusive variable exons and exon selection can be mediated by the splicing regulator Srrm234 in *Drosophila* ^43–47^.

ELAV (Embryonic Lethal Abnormal Visual System)/HU proteins are important neuronal RNA binding proteins, highly conserved and extensively used as neuronal markers ^48–50^. ELAV regulates alternative splicing by binding to AU-rich motifs, which are abundant in introns and untranslated regions ^51^. The *Drosophila* genome has three members of the ELAV family of proteins, while the honeybee genome encodes only one ELAV protein ^30,52,53^. ELAV proteins have prominent roles in regulating synaptic plasticity ^53–57^.

Here, we analysed the combined effects of Thiamethoxam, Carbendazim and Glyphosate on worker bee viability. Further, we determined expression and alternative splicing of *Xbp1, Dscam* and *elav* genes in bees and investigated alternative splicing upon exposure of these commonly used agrochemicals. These experiments could reveal possible indicators of the toxicity of these pesticides. The search for biomarkers and information about the effects of pesticides on the neuronal system of bees is of great importance, aiming to contribute to the characterization of exposure to these xenobiotics at the molecular level.

## RESULTS

### Thiamethoxam toxicity in bees is not enhanced by Carbendazim and Glyphosate

To determine the toxicity of the neonicotinoid thiamthoxam we injected it into worker bees as this is the most accurate way of delivery. We chose a volume of 2 µl for injection based on our experience from injections in *Drosophila* where an estimated 1/10 of the hemolymph volume is well tolerated ^58^. LD50 for Thiamethoxam was between 1 and 10 µM, and 100 µM resulted in complete lethality (Fig 1). Intriguingly, the commonly used fungicide Carbendazim and herbicide Glyphosate at highest water soluble concentrations of 2 mM and 32 mM were not lethal (Fig 1). Furthermore, combining these two compounds with a sub-lethal dose of Thiamethoxam did not enhance its toxicity. (Fig 1). Initially, we assessed viability rates after 24 hours of exposure. We then repeated these injection experiments and determined viability also after 48 hours of exposure to these combinations of agrochemicals and did not observe significant differences between the two time points (Fig 1).

**Figure 1.**
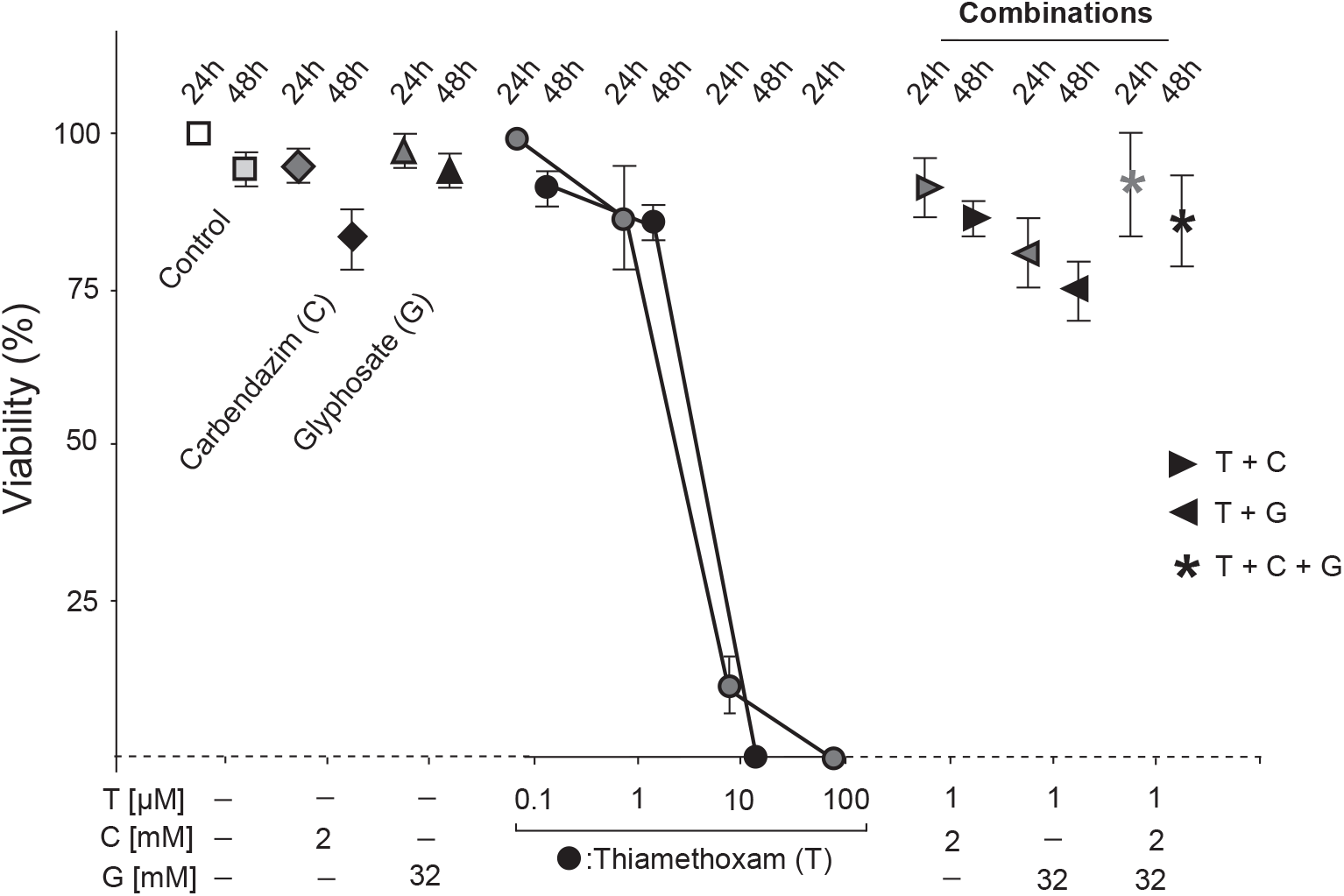
Viability of *Apis mellifera* exposed to xenobiotics. Means with standard errors from three experiments are represented. The percent viability of bees after 24 h and 48 his plotted against the concentration of xenobiotics. Bees were injected with 2 µl of water (Control), Carbendazim (C, 2 mM), Glyphosate (G, 32 mM) and/or Thiamethoxam (T) individually or in combinations at indicated concentrations. The 24 h is indicated by lighter colour and the 48 h time-point by a darker colour.

### *Apis mellifera* alternative splicing of the stress sensor *Xbp1* does not change upon exposure to xenobiotics

Upon cellular stress, the unfolded protein response (UPR) is triggered by non-spliceosomal tRNA-type alternative splicing of the *Xbp1* gene ^31–34, 59^. To confirm that we can induce UPR ^59^, we injected Dithiothreitol (DTT) and followed *Xbp1* splicing by RT-PCR over 24 h at selected time-points (Fig2A-D). Twenty four hours after injection of 2 µl of a 20 mM DTT solution, splicing of *Xbp1* increased about two-fold (Fig 2C and D). In contrast, injection of sub-lethal doses of Thiamethoxam, or the combination of Thiamethoxam with Carbendazim and Glyphosate did not result in apparent changes in the *Xbp1* alternative splicing (Fig 2E).

**Figure 2:**
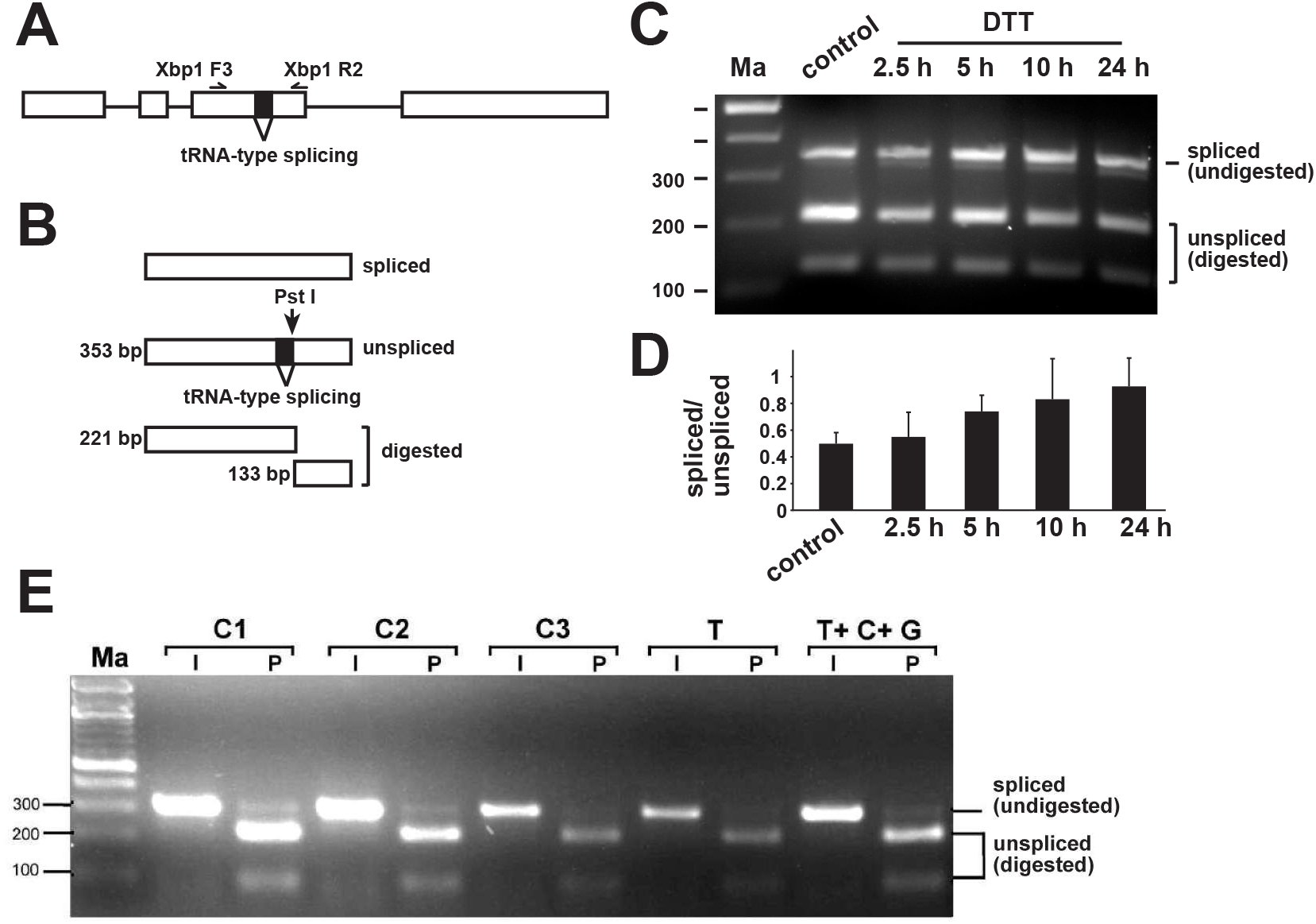
*Apis mellifera Xbp1* non-spliceosomal intron splicing in worker bees is unaffected by thiamethoxan, carbendazim and glyphosate. (A) Gene structure of *Apis mellifera Xbp1* depicting the tRNA-type spliced intron and primers used to analyse its splicing (top). (B) To resolve similar sized spliced and unspliced isoforms, the RT-PCR product was digested with *Pst*I, which only cuts the unspliced RT-PCR product. The size of the smaller fragment for return primer Xbp1 R1 is 91 bp (not shown). (C) Agarose gel showing the alternative splicing pattern of *Xbp1* amplified with primers Xbp1 F3 and R2 by digestion of the RT-PCR product with *Pst*I (P) at different time-points after injection of 2 µl 20 mM DTT. (D) Quantification of the changes in *Xbp1* splicing shown in Fig 2C as mean with the standard error from three replicates of the ration of spliced to unspliced. Only the large fragment of unspliced was used for quantification. (E) Agarose gel showing the alternative splicing pattern of *Xbp1* amplified with primers Xbp1 F3 and R1 by digestion of the RT-PCR product with *Pst*I (P) compared to undigested input (I) in control bees dissected immediately after collection (Control 1), control bees fed with water and sucrose for 24 h (Control 2) and control bees injected with water (Control 3) compared to bees injected with Thiamethoxam (1 µM) and bees injected with a mixture of Thiamethoxam (1 µM, T), Carbendazim (2 mM, C) and Glyphosate (32 mM, G) 24 h prior dissection. Samples were run on a 3% agarose gel. Ma: DNA marker. The undigested PCR product is shown at the bottom.

### *Apis mellifera Dscam* exon 4 alternative splicing does not change during development, in adults and upon exposure to xenobiotics

To examine potentially toxic effects of Thiamethoxam on alternative splicing regulation in bees, we chose to analyse the splicing pattern in one of the most complex genes in arthopods, the *Dscam* gene ^38^. *Dscam* in bees has three variable clusters of mutually exclusive exons which are the exon 4 cluster with 8 annotated variables, the exon 6 cluster with 45 variables and the exon 9 cluster with 17 variables ^60^. We chose the exon 4 cluster because we could separate all variable exons after digestion based on annotated sequences with a combination of restriction enzymes on denaturing polyacrylamide gels, whereby exons 4.1, 4.2, 4.6, 4.7, 4.8 were resolved by Sau3AI, exons 4.3, 4.5 by HaeIII and exon 4.4 by MspI (Fig 3A and B) ^46^. Since the splicing pattern has not been characterized before, we then determined whether all eight annotated exons 4 were present in bees (Fig 3A).

**Figure 3:**
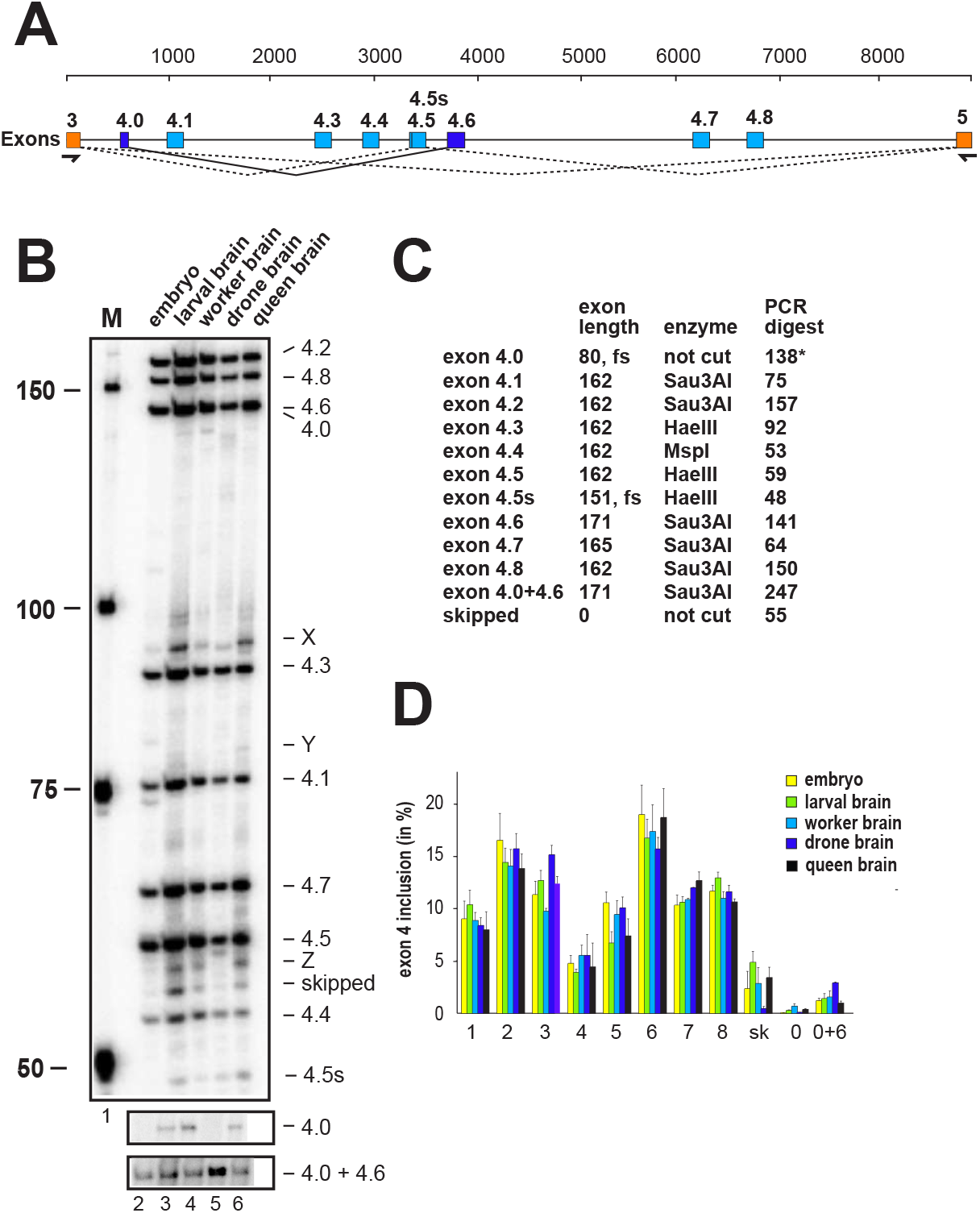
*Apis mellifera Dscam* exon 4 alternative splicing during bee development and between castes. (A) Schematic depiction of *Apis mellifera Dscam* exon 4 variable cluster with primers indicated below orange markes corresponding to constant exons 3 and 5. Variable exons 4 are marked in light blue. Newly discovered exon 4.0 is spliced to exon 4.6 (dark blue). An alternative 5’ splice site discovered in exon 4.5 is indicated as a line. (B) Denaturing polyacrylamide gels (6%) showing the splicing pattern of *Dscam* exon 4 isoform variables on top by digestion of a ^32^P labeled RT-PCR product with a combination of *Hae*III, *Msp*I, and *Sau3*AI restriction enzymes in embryos (line1), larval brains (line 2), worker brains (line 3), drone brains (line 4) and queen brains (line 5). Exon 4.0, that is close to exon 4.6 in length and exon 4.0+4.6 are shown from an undigested control (bottom). (C) Table showing the length of variable exons and their length after restriction digest with indicated restriction enzymes. Exon 4.0 is close to exon 4.6 in length and is shown from na undigested control gel in B. (D) Quantification of inclusion levels of individual exons are shown as means with standard error from three experiments for embryos, larval brains, worker brains, drone brains and queen brains.

Indeed, we could detect all annotated eight exons, but in addition, we also detected an additional exon, termed exon 4.0 (80 nts, CTGTTTAGAA…TACAGACACG), that is mostly spliced to exon 4.6 and a recessed alternative 5’ splice site in exon 4.5. In addition, a number of bands were evident (X-Z, Fig 3B), that could not be further identified using separation and excision of bands on agarose gels for sequencing. In contrast to *Drosophila*, only eight of the 12 exons are present in the bee *Apis mellifera* (Fig 3B and C).

Inclusion levels of variable exons were determined for bee embryos, larval brains, and brains from foragers, drones and queens. The inclusion of annotated exon 4 variants (exons 4.1-4.8) revealed no apparent differences in these five developmental stages (Fig 3D).

After exposure to sub-lethal doses of Thiamethoxam, or the combination of Thiamethoxam with Carbendazim and Glyphosate, no apparent changes in the *Dscam* exon 4 splicing pattern in the brain were detected 5 h, 10 h or 24 h after injection (Fig 4 and Supplemental Fig 2).

**Figure 4:**
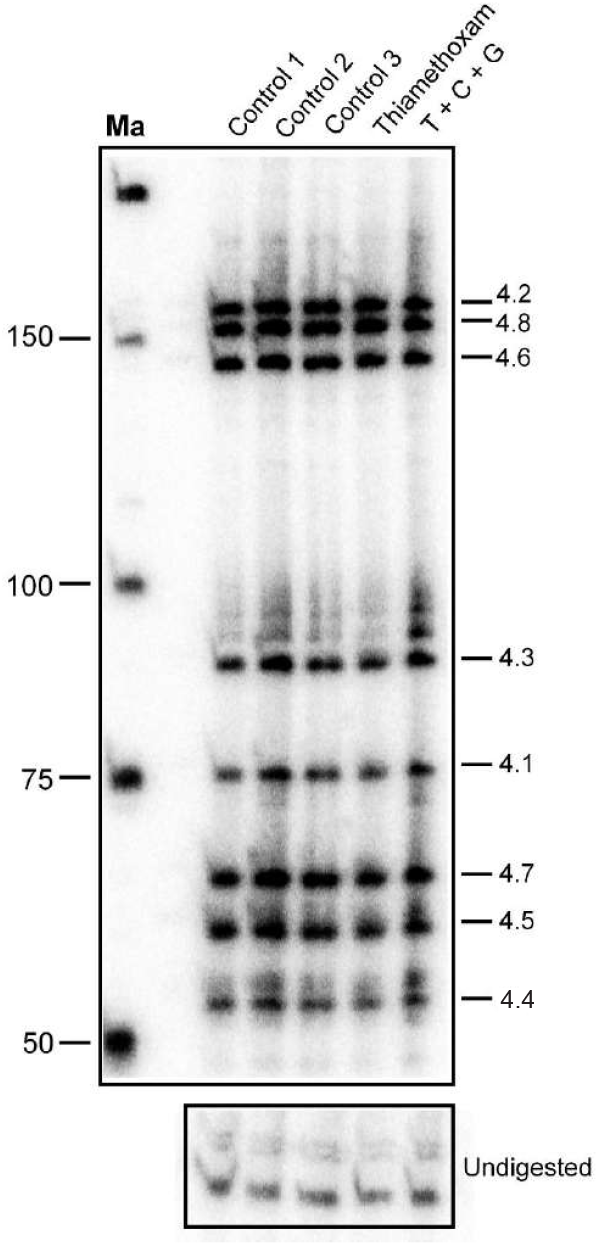
*Apis mellifera Dscam exon 4* alternative splicing in brains of worker bees is unaffected by thiamethoxan, carbendazim and glyphosate. Denaturing polyacrylamide gels (6%) showing the splicing pattern of *Dscam* exon 4 isoform variables on top by digestion of a ^32^P labeled RT-PCR product with a combination of *Hae*III, *Msp*I, and *Sau3*AI restriction enzymes in control bees dissected immediately after collection (Control 1), control bees fed with water and sucrose for 24 h (Control 2) and control bees injected with water (Control 3) compared to bees injected with Thiamethoxam (1 µM) and bees injected with a mixture of Thiamethoxam (1 µM, T), Carbendazim (2 mM, C) and Glyphosate (32 mM, G) 24 h prior dissection. Samples were run on 8% polyacrylamide gel. Ma: DNA marker. The undigested PCR product is shown at the bottom.

### *Apis mellifera elav* alternative splicing does not change upon exposure to xenobiotics

Next, we examined alternative splicing of *elav* in honeybee workers upon exposure to xenobiotics. To determine alternative splice forms, 5’ and 3’ ^32^P labelled PCR products covering the variable region were digested with *Kpn*I and *Fok*I restriction enzymes, respectively, and resolved by denaturing polyacrylamide gels (Fig 5A-C). After exposure for 24 h to sub-lethal doses of Thiamethoxam, or the combination of Thiamethoxam with Carbendazim and Glyphosate, no apparent changes were detected in the *elav* splicing pattern in the brain (Fig 5B and 5C).

**Figure 5:**
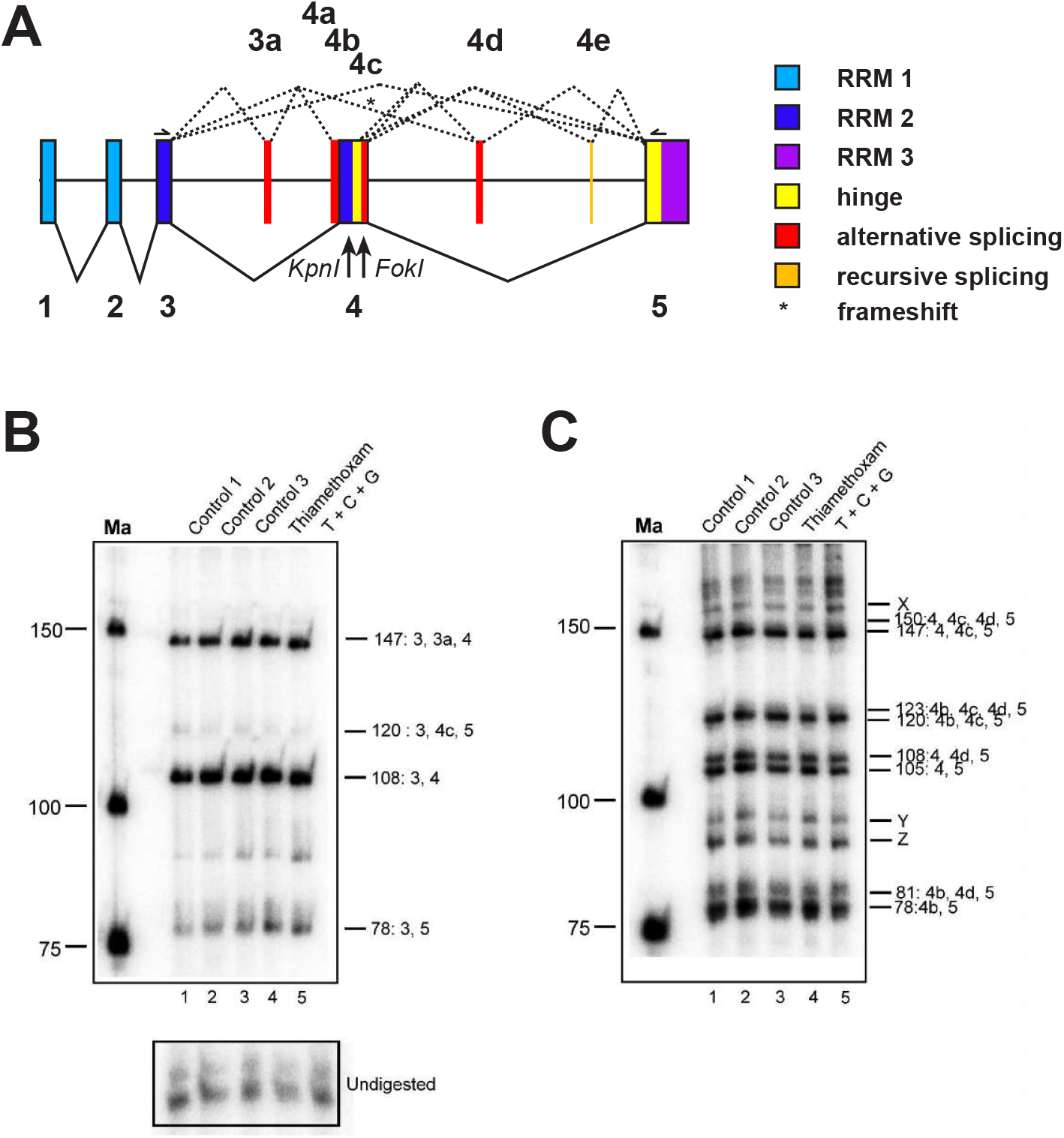
*Apis mellifera elav* alternative splicing in brains of worker bees is unaffected by thiamethoxan, carbendazim and glyphosate. (A) Gene structure of *Apis mellifera elav* depicting color-coded functional protein domains with constant exons (1-5, bottom, solid lines) and alternative splicing exons (3a and 4a-d, top, dashed lines). RNA Recognition Motiv 1 (RRM1): light blue, RRM2: dark blue, RRM3: purple, hinge region: red and alternatively spliced parts in red. *Kpn*I and *Fok*I restriction sites used to separate isoforms are indicated below the gene model. An asterisk indicates isoforms that encode truncated proteins by introducing a frameshift. (B, C) Denaturing polyacrylamide gels (6%) showing the alternative splicing pattern of *elav* by digestion of a 5’ (B) or 3’ (C) ^32^P labeled RT-PCR product with *Kpn*I (B) and *Fok*I (C) in control bees dissected immediately after collection (Control 1), control bees fed with water and sucrose for 24 h (Control 2) and control bees injected with water (Control 3) compared to bees injected with Thiamethoxam (1 µM) and bees injected with a mixture of Thiamethoxam (1 µM, T), Carbendazim (2 mM, C) and Glyphosate (32 mM, G) 24 h prior dissection. Samples were run on 6% polyacrylamide gel. Ma: DNA marker. The undigested PCR product is shown at the bottom.

## DISCUSSION

Like other insects, honeybees are exposed to complex combinations of agrochemicals present at potentially harmful concentrations in agricultural landscapes ^61^. Honeybees display marked variation in sensitivity to different insecticides ^62,63^, which may be influenced by different types of pesticide-pesticide cocktails ^61,24,64^.

Here we investigated for the first time whether alternative splicing of important stress and neuronal genes is activated in the nervous system of the honeybee following acute exposure to xenobiotics. Injection into the hemolymph of forager bees ensured accurate and identical application of the tested pesticides. This is an initial approach to explore molecular effects of compounds on individual forager bees before evaluating exposure outcomes at the colony-level in cost- and labor-intensive field-experiments^65^.

Within a 48 hour window after acute exposure we did not observe any synergistic or additive effects on survival rates when sub-lethal doses of either glyphosate or carbendazim, or both, were administered along with the neonicotinoid Thiamethoxam. Phenotypic outcomes of these particular combinations of insecticide, herbicide and fungicide have not been reported previously. As we only examined the viability of individual forager bees following short-term exposure we cannot rule out other, more subtle phenotypic effects nor predict what the synergistic impacts of chronic exposure to these pesticide combinations would be on individual bees or on a hive level. For example, gene expression alterations have been detected following 48 hour and 72 hour Thiamethoxam exposures at concentrations which are similar to the sub-lethal dose tested here, however these changes after such prolonged time are likely indirect ^66^.

Alternative splicing is a mechanism by which the exons are spliced in different ways to generate multiple transcripts from one mRNA precursor. This process contributes to protein diversity by generating different types of proteins and has been recognised as a rapid cellular mechanism in response to environmental perturbation ^27^. From our experience with *Drosophila*, alternative splicing differences are most pronounced 24 hours after xenobiotic exposure ^30^;; for this reason we chose this time point to investigate RNA splicing in honeybees.

Alternative splicing is further thought to provide means for adaption to environmental changes, but given the complexity of the splicing process involving hundreds of proteins alternative splicing likely is also susceptible to interference by xenobiotics ^29,30^. Since neonicotinoids show neurotoxic features in bees, we reasoned that alternative splicing of *elav* might be altered when bees are exposed to Thiamethoxam alone, or in combination with glyphosate and carbendazim. Therefore, we analyzed the inclusion levels of *elav* variable exons through a novel method ^46^. Our results revealed minimal changes in *elav* splicing in the presence of the abiotic pesticide stressors when compared to the control groups.

Similar to *elav*, the same pesticide dosages and exposure conditions revealed no significant changes in splicing patterns of *Dscam* exon 4. This lack of alternative splicing changes was unexpected as an enormous *Dscam* diversity is generated by mutually exclusive splicing in the *Drosophila* nervous and immune systems ^38^.

The third gene we investigated for alternative splicing was *Xbp1*. *Xbp1* is involved in the unfolded protein response (UPR), which is activated during stress conditions ^67^. Johnston and colleagues had previously reported a robust UPR activation in the honeybee in response to multiple known stressors, including tunicamycin, a protein glycosylation inhibitor and DTT ^59^. Based on these findings we reasoned that *Xbp1* might serve as a key molecular component in mediating individual and combined effects of environmental stressors in honeybees. However, we did not detect changes in the characteristic tRNA-type cytoplasmic splicing event that processes *Xbp1* transcripts in response to cellular accumulation of unfolded proteins. Although our study is limited in that we evaluated potential alternative splicing events in transcripts of only three genes, namely *Xbp1, Dscam* and *elav*, these genes are important representatives of regulators of a stress-response, neuronal wiring, the immune response and synaptogenesis important for behavioural performance during foraging. For *Xbp1, Dscam* and *elav* acute exposure in foraging honeybees does not trigger evolutionarily conserved stress-related alternative splicing processes in brain derived RNA. It remains to be shown by large-scale analysis of alternative splicing from whole genome RNA sequencing whether the lack of activation of this RNA processing mechanism is a general feature in honeybees. If alternative splicing remains underused as an adaptive response, this would leave the western honeybee more vulnerable to man-made environmental stresses.

## MATERIALS AND METHODS

### Toxicity assays

For developmental expression studies, bees (*Apis mellifera*) of different castes and developmental stages were taken from the experimental apiary of the University campus in Toulouse (France), and cold-anesthetized by placing them on ice before dissection. Forager bees for toxicity assays were collected from colonies of the Winterbourne Garden of the University of Birmingham (UK) and kept in small round cages used for food storage (500 ml) with holes for air circulation. Bees were fed daily from a sucrose (1:1) filled 2 ml Eppendorf tube with small holes inserted into the lid of the container through a hole. Water was provided by a paper tissue saturated daily with water (Evian). For each experimental group 30 bees were collected (replicates with ten individuals in each group). To ensure that laboratory conditions were not stressful for bees, three groups of 10-12 bees were used. Bees from control group 1 were dissected immediately after collection, and their brains extracted. Bees from control group 2 were fed and dissected after 24 hours. Bees from control group 3 were injected with water into abdomen. Thiamethoxam (Sigma-Aldrich, #37942, PESTANAL analytical standard), Carbendazim (Sigma-Aldrich, #45368, PESTANAL analytical standard) and Glyphosate (Round-up, Bayer) were diluted in water at their maximal soluble concentration and then the minimum lethal dose was determined by injection. Two µl Dithiothreitol (DTT) was injected as a 20 mM solution in water. For injections bees were cold-anaesthetised by placing them on ice and then put on a custom made metal pad connected to circulating ice-water driven by an aquarium pump to keep them anaesthetised during injections. Injections were done with a 10 µl Hamilton syringe and each cold-anaesthetised bee was injected with 2 µl of individual xenobiotics are a mixture thereof with indicated concentrations into the left side of the abdomen between abdominal segment 2 and 3 (Supplemental Figure 1). After injections, bees were kept in a humidified incubator at 32°C and viability was scored after 24 hours.

### RNA extraction, reverse transcription (RT) and polymerase chain reaction (PCR) and analysis of alternative splicing

RNA extraction was done using Tri-reagent (SIGMA) and reverse transcription was done with Superscript II (Invitrogen) as previously described ^68^ using primer AM Dscam 13R2 (GCCGAGAGTCCTGCGCCGATTCCATTCACAG, 1 pmol/ 20 µl reaction) in combination with an oligo dT primer. 1-2 µl of the RT-PCR mix was used in 50 µl PCR reaction using Fermentas Taq (Thermo Scientific) according to the manufacturer’s instructions. *Xbp1* was amplified with primers Xbp1 AM F3 (GAAGAAACTGTTCGAAGGTTAAGGGAAC) and Xbp1 AM R2 (GTTCGATATAATCATCTCCTTGGAG) or Xbp1 AM R1 (TCAAGAGGAAGTAGATGGTCAGAA). For Pst I digestion 40 u of the enzyme was added after PCR to the mix and digested for 1 h. PCR products were then analysed on ethidium bromide stained 3% agarose gels. To amplify the *Dscam* exon 4 cluster, PCR was performed using primers AM Dscam 3F1 (AGTTCACAGCCGAGATGTTAGCGTGAGAGC) and AM Dscam 5R1 (GGAAGGCAGTACCAAGTATTTTC) for 37 cycles with 1 µl of cDNA. New variables of *Dscam* exon 4 were gel purified and determined by sequencing (exon 4.0 and 4.0+4.6) or by the annotated sequence (Exon 4.5 recessed alternative 5’ splice site). *Apis elav* was amplified with primers elav AM F2 (GTCGCGGATACTTTGCGACAACATCAC) and elav AM R2 (CCCGGGTAGCATCGAGTTTGCCAATAGATC). For the analysis of *Dscam* and *elav* alternative splicing primers were labeled with ^32^P gamma-ATP (6000 Ci/ mmol, 25 mM, Perkin Elmer) with PNK to saturation and diluted as appropriate ^69^. From a standard PCR reaction with a ^32^P labelled forward primer, 10–20% were sequentially digested with a mix of restriction enzymes according to the manufacturer’s instructions (NEB) ^69^. PCR reaction and restriction digests were phenol/CHCl_3_ extracted, ethanol precipitated in the presence of glycogen (Roche) and analyzed on standard 6% sequencing type denaturing polyacrylamide gels. After exposure to a phosphoimager (BioRad), individual bands were quantified using ImageQuant (BioRad) and inclusion levels for individual variable exons were calculated from the summed up total of all variables. Statistical analysis was done by one-way ANOVA followed by Tukey– Kramer post-hoc analysis using Graphpad prism. Percent inclusion levels were calculated from the total sum of variables as described ^46^.

## AUTHORS’ CONTRIBUTION

R.S. T.C.R, and M.S conceptualized the project. P.D., P.U. performed the experiments, T.C.R, O.M., J.M.D, R.S. and M.S. supervised experiments and analyzed data, R.S. and M.S. wrote the manuscript with help from P.D., P.U. and J.M.D.

## ACKNOWLEDGMENTS

We thank Noel Parker and the Winterbourne garden for bees, Noel Parker for bee suits, V. Soller-Haussmann for help with bee collections, and L. Hotier for collecting bees at the Toulouse apiary. For this work we acknowledge funding from the Foundation for Research Support of the State of São Paulo, FAPESP (2012 / 13370-8;; 2014 / 23197-7), the Biotechnology and Biological Sciences Research Council (BBSRC), the Nottingham Birmingham Fund, the Genetics and the Biochemical Society, and the Sukran Sinan Memory Fund.

## DATA AVAILABILITY

The datasets generated during and/or analysed during the current study are available from the corresponding author on reasonable request.

